# The Effects of Social Stress on Memory: If, How and When

**DOI:** 10.1101/2020.10.01.322271

**Authors:** Elizabeth McManus, Deborah Talmi, Hamied Haroon, Nils Muhlert

## Abstract

Physical stress, such as from the cold-pressor test, has been robustly associated with altered memory retrieval, but it is not yet clear whether the same happens following psychosocial stress. Studies using psychosocial stressors report mixed effects on memory, leading to uncertainty about the common cognitive impact of both forms of stress. The current study uses a stepped replication design, with four near-identical experiments, each differing by a single critical factor. In three experiments we induced psychosocial stress after participants encoded word stimuli, then assessed retrieval after a prolonged delay. These experiments found no group level influence of postencoding stress on recognition of neutral words or cued recall of word-pairs, but a small effect on recollection of semantically-related words. There was, however, some indication of positive relationships within the stress group between measures of stress (cortisol in experiment 1 and self-reported-anxiety in experiment 3) and recollection of single word stimuli. In the fourth experiment, we found that psychosocial stress immediately before retrieval did not influence word recognition. Overall, our findings demonstrate that psychosocial stress has a typically modest impact on memory, lower than previously claimed, but that individual differences in stress responsivity, particularly for tasks that tap recollection, may help to explain variability in previous findings.

## 1. Introduction

It is commonly believed that at times of great stress our behaviour changes. Typically this is considered to be changes in our ability to make decisions, but acute stress might also affect our ability to retrieve known information or learn new information (Lupien & Lepage, 2001). The direction of the effect of acute stress on behaviour is thought to depend on when it occurs: if stress occurs close to the time of learning it enhances memory, whereas stress immediately before retrieval impairs memory (Shields, Sazma, McCullough, & Yonelinas, 2017; Wolf, 2017). Yet this understanding has been based mainly on animal models, or on findings that do not discriminate between physical and psychosocial stressors. It is not clear to what extent the same conclusion characterises the effect of stressors in everyday life, where a substantial proportion of the stress humans experience is psychosocial. In the present experiments, we use a careful, stepped experimental design, to determine what effect, if any, psychosocial stress has on human memory.

Even within the psychosocial stress literature, there is considerable variability of findings about the effect of stress as a function of its induction time relative to the learning episode (Beckner, Tucker, Delville, & Mohr, 2006; Corbett, Weinberg, & Duarte, 2017; Sheldon, Chu, Nitschke, Pruessner, & Bartz, 2018; Tollenaar, Elzinga, Spinhoven, & Everaerd, 2009). In part, this may be due to inter-individual variability in stress response, but also to the heterogeneous nature of memory itself, as this construct has been operationalised in many different ways in that literature. Free recall tasks are completed by searching and actively recalling previously learned stimuli, whereas recognition memory is typically assessed using old/new identification tasks. Exploration of the processes that underlie item recognition can also be carried out using remember-know tasks: these assess levels of recollection and familiarity (e.g. Migo, Mayes, & Montaldi, 2012). In remember-know tasks, participants are asked if they have a sense that test stimuli have been encountered before (responding ‘know’ to represent familiarity) or if they have a vivid memory of some other specific details from the previous encounter with the stimuli (responding ‘remember’ to represent recollection). It is also possible to test recognition and recall jointly, in paired-associates tasks. For example, participants could initially be asked to identify if a stimulus has been encountered before (item recognition), and then asked to recall what other stimulus it was paired with (cued recall) (Old & Naveh-Benjamin, 2008).

Familiarity, recollection and free recall, may differentially recruit the neural circuitry implicated in stress responses, which could potentially lead to variable effects of stress on memory (Dedovic, Duchesne, Andrews, Engert, & Pruessner, 2009; McEwen, 2007). Previously, it has been proposed that only hippocampally-dependent memory could be influenced by stress (McEwen, 2007). This amounts to the suggestion that recollection, which is often believed to be uniquely based on the hippocampus (Montaldi & Mayes, 2010), will be particularly strongly affected. However, more recent evidence suggests that stress may also influence other, less hippocampally-dependent memory processes, including familiarity (Li, Weerda, Guenzel, Wolf, & Thiel, 2013; McCullough & Yonelinas, 2013; Wolf, 2017).

In addition to the memory test, a key factor that may influence the impact of stress on memory is the valence of stimuli. Emotion, whether external to or intrinsic within stimuli, is thought to enhance memory relative to neutral materials (Ponzio & Mather, 2014; Sharot & Yonelinas, 2008). This suggestion can also extend to stress as in general, stress after learning is thought to enhances memory for emotional stimuli relative to neutral stimuli (LaBar & Cabeza, 2006). This finding supports the modulation model, in which emotional memories recruit additional activity in the amygdala, following secretion of noradrenaline, cortisol or other related hormones. This amygdala activity, in turn, aids consolidation, rendering emotional memories less vulnerable to forgetting (McGaugh, 2004). The same increase in amygdala activation is also thought to occur in response to stress and the related increase in secretion of the same noradrenergic and other stress-related hormones, such as cortisol. The combination of both emotional stimuli and increased levels of stress could have additive influence on amygdala activation and ensuing memory enhancement. In practice however, both enhanced and impaired memory for positive and negative emotional materials have been reported after stress (Cornelisse, van Stegeren, & Joëls, 2011; Elzinga, Bakker, & Bremner, 2005; Kamp, Endemann, Domes, & Mecklinger, 2019; Kuhlmann, Piel, & Wolf, 2005; Preuß & Wolf, 2009; Wolf, 2012), alongside null findings (Schoofs & Wolf, 2009; Tollenaar et al., 2009). This heterogeneous picture may reflect differences in the population studied, or in the memory tests the studies employed.

In order to understand when and how stress influences memory, more consistency in study design may be needed to identify the specific combination of crucial factors. A current problem within the literature is the large number of different methods of assessment for memory and the low number of direct replications of such studies. The diversity of methods makes it hard to interpret exactly which factor results in stress-related memory alterations. Meta-analyses have successfully examined the extent to which different factors contribute to the overall effect of stress on memory, but existing meta-analyses (Shields et al., 2017; Wolf, 2009) have not distinguished between physical and psychosocial stress. There is good reason to believe that their effect would differ. For instance, the neural networks activated differ between physiological and psychosocial stress (e.g. Kogler et al., 2015), and the pattern of physiological stress response is known to depend on the specific nature of the psychosocial background (Chida & Hamer, 2008).

In the current study we attempt to overcome these issues by conducting four experiments of near-identical design, each manipulating one crucial factor to allow for a more thorough, stepped approach. Through this we hope to gain a clearer understanding of how and when stress influences memory. Each experiment uses the same sample size, the same experimental stress test and the same active control group. The first three experiments all induce psychosocial stress immediately after learning and have a delay period of 24 hours before examining long term memory. These three experiments differ only in the memory task used. The first experiment assessed recognition of neutral words using an old/new recognition memory test, and also used a remember/know task to assess recollection and familiarity-based recognition memory. The second and third experiments closely replicated the first, but with some differences to address additional questions. The stimuli used in the second experiment were neutral word pairs, not single words. The first step of the memory task replicated the old/new item recognition test (carried out by presenting one word from each pair). This was followed up using a cued-recall test for the second word in each pair. For the third experiment, in addition to neutral words, an additional set of categorised neutral words and a set of emotionally-negative words were also used as stimuli. This experiment used the same old/new recognition and remember/know tasks as experiment one The fourth experiment uses an identical memory procedure to the first, but induced stress immediately before retrieval instead of after encoding. The overall design allows for clearer insight into the effects of stress at different types and stages of memory.

Salivary cortisol, a measure of the physiological response to stress, and self-reports of state anxiety were measured at multiple timepoints while participants underwent the Trier Social Stress Test (TSST; Kirschbaum, Pirke, & Hellhammer, 1993), a commonly used lab-based experimental psychosocial stress paradigm. If this stress manipulation is effective, we predict significantly higher scores for the stress group relative to controls on both these measures. Personality factors, specifically trait stress reactivity scores, were also measured to better identify potential stress responders and non-responders within the stress and control groups.

In line with previous meta-analyses, we expected to see memory enhancements in the stress group relative to controls in the three experiments using a post-encoding stress design. Beyond enhanced recognition memory for the stress group relative to controls during the old/new recognition task, we predict that there will be specific enhancement in recollection ability across all these experiments as this process is thought to be mediated by the hippocampus, a region also closely related specifically to social stress response. For the fourth study, inducing stress immediately before retrieval, based on evidence from other literatures, we would expect to see impairment in memory for the stress group relative to controls. Again, we expect this impairment to be more prominent when considering recollection, but also for recognition accuracy. Last, beyond group effects of stress, we expected to see the greatest impact of stress on memory in participants who experience the greatest levels of stress. We predicted there will be significant correlations between levels of stress, as measured both by self-reports and salivary cortisol, and memory scores, within the stress group. In experiments 1,2 and 3, we predict these correlations will be positive, demonstrating increased levels of stress leads to increased memory enhancements, whereas in experiment 4, we predict this correlation to be negative, indicating further impairments in memory due to stress.

## 2. Methods

### 2.1 Participants

An a-priori power calculation was carried out using G-power software to determine appropriate sample sizes to compare memory scores between groups. It was determined that in order to detect a medium effect size (Cohen’s d = 0.5; (Cohen, 1988)) with a power of .8 (alpha = .05), 18 participants per group would be required. The medium effect size was estimated based on similar previous research (Nater et al., 2007). Thirty-six healthy undergraduate participants were recruited for each of the four studies. Recruitment for each experiment occurred separately, but participants who had taken part in one experiment were excluded from subsequent ones. Demographic data for all four experiments are shown in Table 1. Stress and control groups did not differ in terms of age or sex.

**Table 1.** Demographics data for participants in stress and control groups for all four studies. Age data presented as mean and standard deviation. Number of saliva samples missing for analysis in each experiment is also reported.

|  | Stress |  |  | Control |  |  |
| --- | --- | --- | --- | --- | --- | --- |
|  | Age | Sex<br>(Male/Female) | Missing<br>Saliva Data | Age | Sex<br>(Male/Female) | Missing<br>Saliva Data |
| Experiment 1 | 20.53<br>(2.07) | 7 / 11 | 2 | 20.80<br>(2.37) | 5 / 13 | 1 |
| Experiment 2 | 19.63<br>(1.09) | 5 / 13 | 4 | 21.69<br>(2.87) | 4 / 14 | 4 |
| Experiment 3 | 21.89<br>(3.38) | 3 / 15 | 3 | 21.28<br>(3.06) | 8 / 10 | 3 |
| Experiment 4 | 19.31<br>(1.35) | 7 / 11 | 0 | 20.56<br>(2.19) | 4 / 14 | 2 |

Participants were recruited from the University of Manchester and received either course credits or £9 cash for taking part. Inclusion criteria required all participants to be between the ages of 18-30 and have no history of neurological or psychiatric disorders. Allocation to stress and control groups were pseudo-randomised. All stress sessions occurred in the afternoon (12:00-18:00) when baseline cortisol is at its lowest (Fries, Dettenborn, & Kirschbaum, 2009). Participants were asked not to eat, drink or smoke for two hours prior to the stress session. All experiments were conducted in accordance with the declaration of Helsinki. This study was approved by the University of Manchester research ethics committee.

### 2.2 Materials

The words in all experiments were taken from the Emote database (Grühn, 2016). We used 60 target words in experiments 1, 3 and 4, 60 word-pairs in experiment 2, and 60 foil-words (or word pairs) in all experiments. All participants learned the same target words during the learning phase. Two retrieval lists were constructed, which included an even number of targets and foils, and allocated to the immediate and delayed retrieval tasks, counterbalanced across participants. Learnt words and foils did not statistically differ on any dimension, including word length, frequency, imagery and meaningfulness (see Table 2). Additionally, across experiments, words did not significant differ on any unexpected dimension. When comparing experiment 3 to other experiments, words did differ significantly for valence and arousal, however this is to be expected as experiment 3 used emotionally-negative stimuli whereas other experiments did not. Experiments 1, 2 and 4 all used neutral words, each scoring within one standard deviation (SDs) above or below the mean scores for valence and arousal. These values also did not differ between learnt and foil words. For experiment 2, initial pilot testing established that the words in each pair were not semantically related. For this purpose, participants were asked to rate on a 1(not at all)-5(very related) scale how semantically related the word pairs were. All pairs scored on low on average (<2), suggesting that pairs were not strongly interrelated.

**Table 2.**
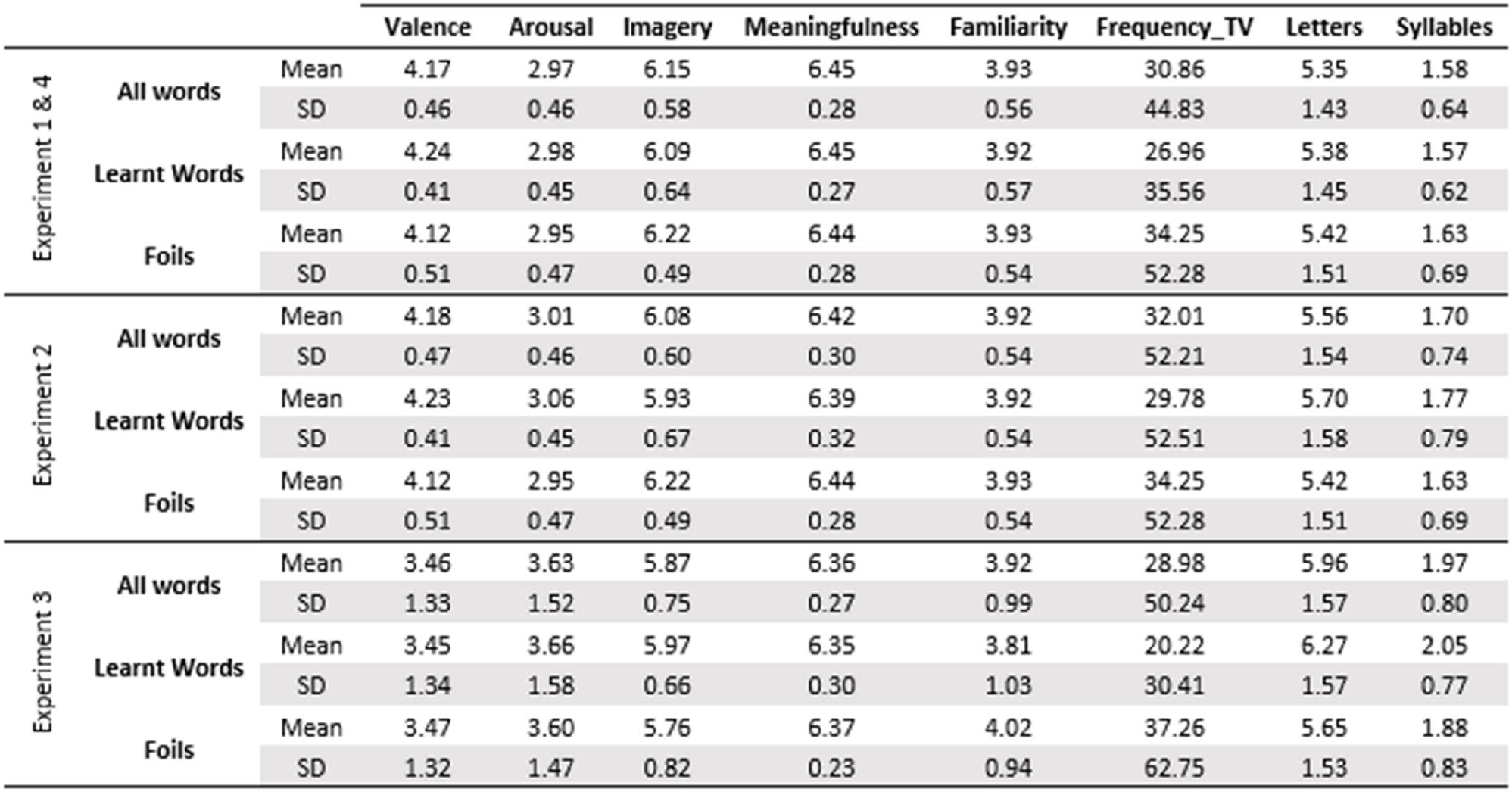
Word stimuli mean and SD for each dimension of interest across all four experiments. Values given for all words in addition to learnt and foil words separately.

Experiment 3 used three types of lists – 20 random-neutral words, 20 related-neutral words, and 20 negative-emotional words. Negative-emotional words were defined as having very low scores for valence, more than one SD different from the mean, and very high scores for arousal, more than one SD above the mean. Within each list type, learnt and foil words did not statistically differ on any dimension (see Table 3). Initial piloting of semantically-related stimuli established that all words belonged to the category of “food and drink”: For this purpose, pilot participants rated 60 words on a scale from 1-10 for their belonging to the semantic category food and drink. We used 20 words, rated above 7, to construct the semantically-related list, and words rated below a 4 for the random list. Significant differences in latent semantic analysis scores between list types suggest that words in the related-neutral list were more semantically related than words in emotionally-negative (*t*(65.917) = 5.28, *p* < .001) and random-neutral lists (*t*(56.1) = 9.56, *p* < .001). As expected, emotionally-negative words were also more semantically related than random-neutral words (*t*(78) = 5.48, p < .001).

**Table 3.**
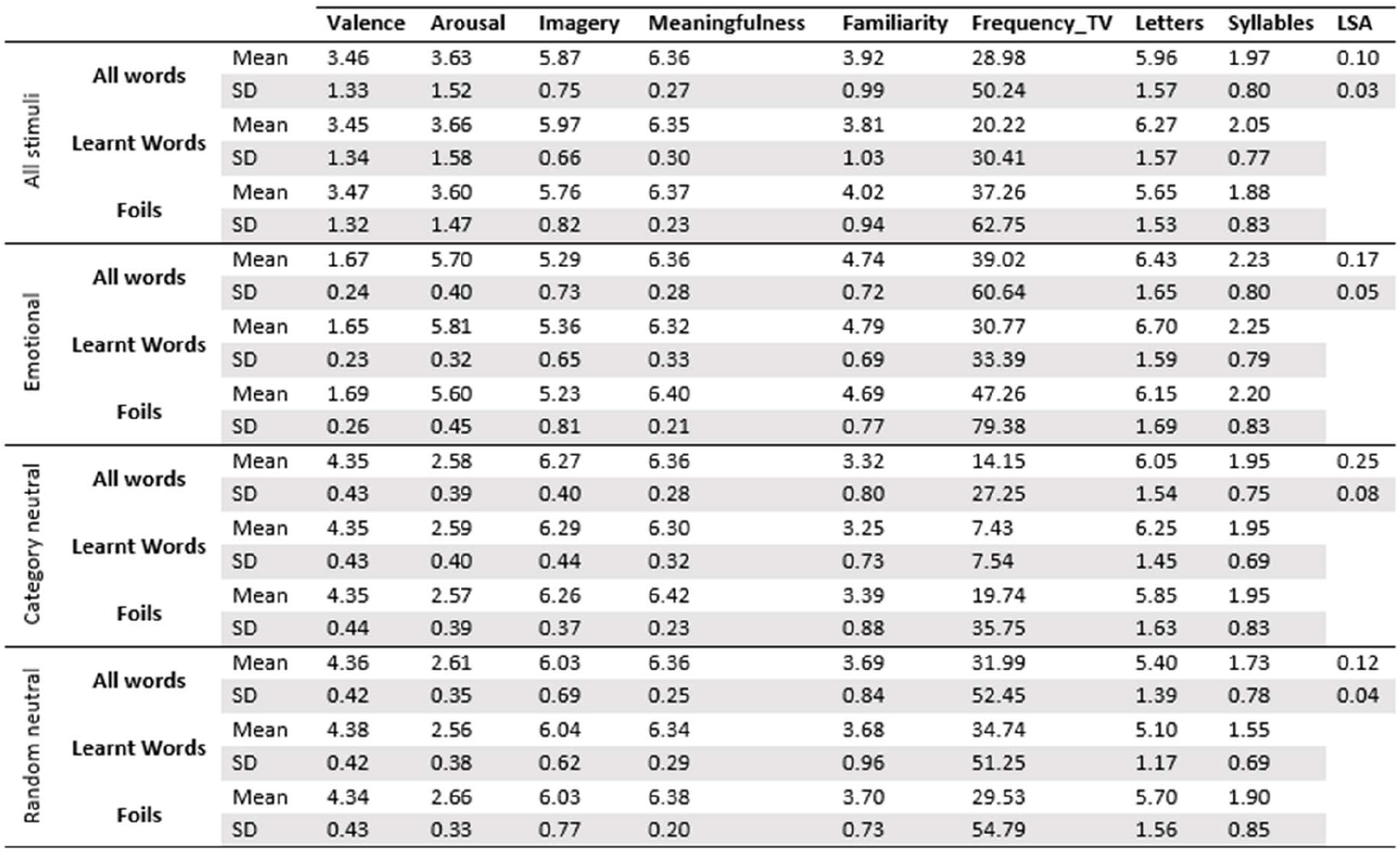
Experiment 3 stimuli word stimuli mean and SD dimension values separated by list type. Values given for all words in each list in addition to learnt and foil words separately.

All participants completed visual analogue scales (VAS) five times throughout the stress session. In addition, participants completed the Perceived Stress Reactivity Scale (PSRS) (Schlotz, Yim, Zoccola, Jansen, & Schulz, 2011). This trait measure of stress reactivity was administered during the non-stressful session to avoid any influence of acute stress.

### 2.3 Memory Tasks

#### 2.3.1 Experiment 1 & 4: Neutral Item Recognition

The participant was shown a list of 60 neutral words presented one at a time on a computer screen. They were instructed to watch the words and remember as many as possible for a later memory task. After the brief distractor task, participants were instructed to complete the recognition task. They were presented with a set of single words but would be asked if each word was “old” and had been shown in the learning list, or “new” and hadn’t been presented to them before. If participants answered “old” they were subsequently asked “do you remember or know you saw this word?”. Remember and know responses were carefully defined to participants before the start of the task to ensure they knew the distinction between the two. The same recognition task procedure was repeated for the delayed recognition task.

#### 2.3.2 Experiment 2: Associative Memory with Cued Pair Recall

The participants were presented with 60 pairs of neutral words, one pair at a time. As with previous experiments, participants were told to remember as many of the word pairs as possible for a later memory task. After the distractor task participants were shown one word at a time and asked to indicate if the word was “old” (they had seen it in the learning list), or “new” (it had not been seen before). When participants answered “old” they were prompted with a further question “what other word appeared with this word” requiring them to recall the other half of the word pair. Only one word from each pair was presented to participants across the two recognition tasks. Target words were counterbalanced so equal numbers of words tested appeared on the left and on the right side of the pair.

#### 2.3.3 Experiment 3: Emotional Item Recognition

The memory task for this experiment follows the same procedure as experiment 1, but used 20 negative-emotional, 20 random-neutral and 20 related-neutral (food) words. These words were presented as part of a mixed list and the distinction between word types was never explicitly discussed with participants, although they were warned in the PIS regarding the possibility of negative-emotional words. Equal numbers of each types of words were used in the immediate and delayed tasks and in the list of foils.

### 2.4 Procedure

The study took place across two sessions, separated by 1 day. Stress occurred on day 1 for experiments 1, 2 and 3, but on day 2 for experiment 4. The study took place in two rooms, A and B. Participants were led between rooms A and B during the session by the researcher. TSST panel members remained in room B on the day the stress task occurred and were not present on the other day.

Five times throughout the testing session including the stress task, participants were asked to add a mark to a 10cm line that best described their current feelings of anxiety, from “not at all anxious (0)” to “extremely anxious (10)”. The five time points were: (1) immediately after baseline rest period (2) after preparation for the Trier (3) after the Trier speech task (4) after the Trier maths task (5) immediately before study debrief. At time points 1, 4 and 5 passive drool saliva samples were also collected. Stress levels were not measured on the other day of testing.

#### 2.4.1 Post-encoding stress experiments 1, 2 and 3

The timeline for all post-encoding stress experiments can be seen in Figure 1a. On day 1 the participant initially provided informed consent before being left alone in room A for a 10-minute rest period. Afterwards, the participant provided the first anxiety rating and saliva sample, answered some general health questions, and then completed the learning task. For this, participants were presented with a list of 60 words (or pairs of words in Experiment 2) one at a time, with each word (or word pair) remaining on screen for 3 seconds followed by a 2 second fixation cross. Before the task began, participants were asked to try to remember as many of the words as possible for a subsequent memory task. Immediately following the learning task, participants were presented with a distractor task, requiring them to read aloud a passage of text for 30 seconds. Following the distractor task, the participants completed the relevant immediate recognition task, details of which changed between experiments, so are discussed individually.

**Figure 1.**
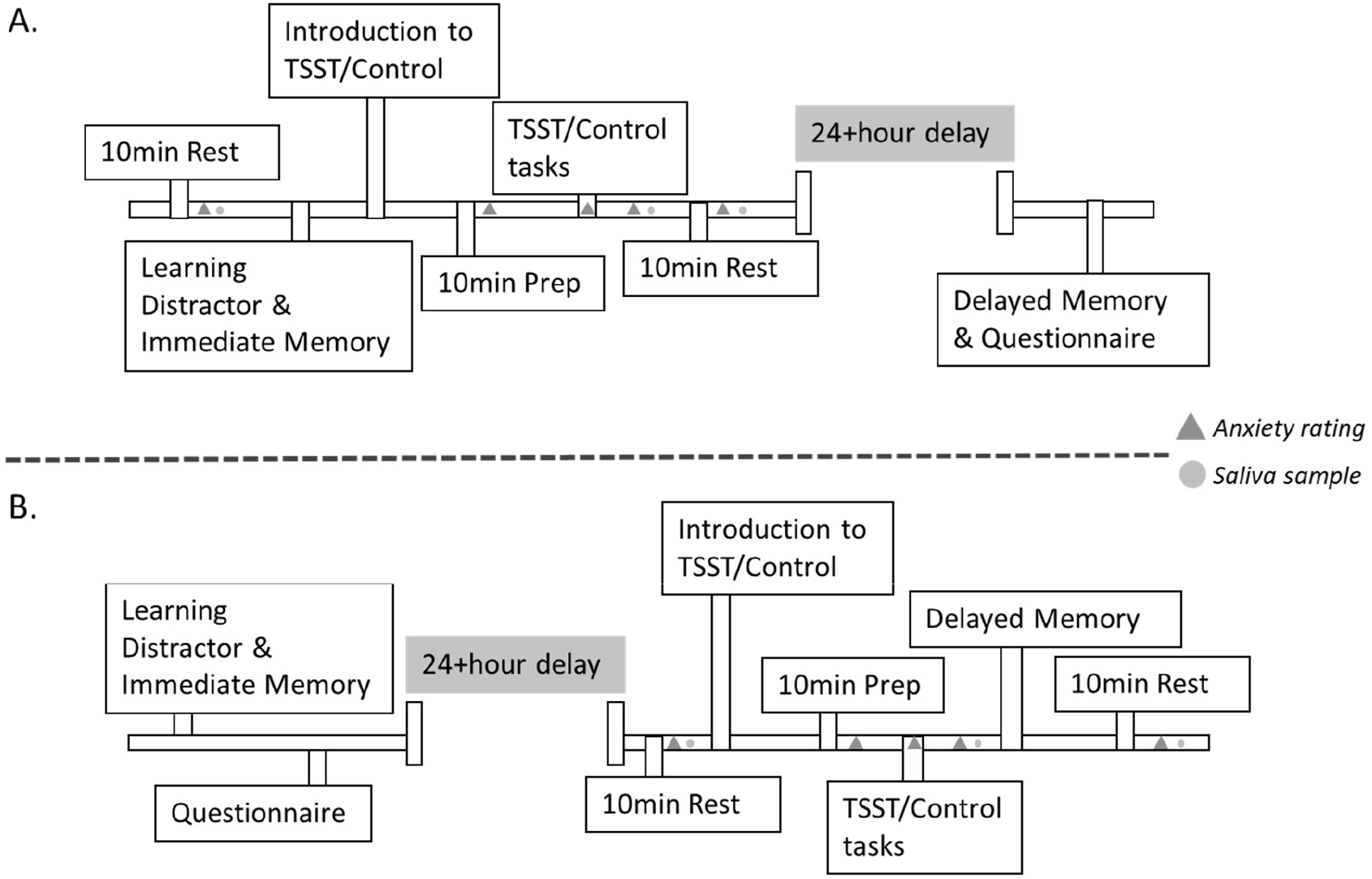
Timelines of tasks. Panel A shows the timeline for Experiments 1, 2 and 3, which administered stress in the same session as the learning, after the immediate retrieval test. Panel B shows the timeline for Experiment 4, which administered stress just before retrieval.

The participant was taken to room B, where they were introduced to the speech element of the TSST or control task. The stress group were informed they would be required to give a 5-minute speech for a job interview in front of the video camera and panel of two socially rejecting judges and that they would have 10 minutes to prepare for this task. The control group were told they would have a 5-minute conversation with the panellists about a topic of their own choosing and were also given 10 minutes to prepare. Both groups were informed that a second task would follow once the first part was completed. Pen and paper were provided to allow participants to make notes during the preparation phase back in room A, but they were informed they could not take these notes into the task with them. After the preparation period in room A, participants provided a second anxiety rating before returning to room B (for the Trier/Control task).

In the stress condition, the two panellists remained neutral and did not respond to the participant during the speech task. If the participant stopped talking, the panellists offered a prompt or follow up question (e.g. Other than your academic achievements, what qualifies you for this role?) only after a 20second period of silence. In the control task, panellists allowed the participant to lead the conversation but asked friendly questions and showed interest and encouragement throughout. Upon completion of this part, participants completed a third anxiety rating using VAS. All participants were then informed that they were now to complete a mental arithmetic task. For the stress group this task involved counting backwards from 1687 in steps of 13 as fast and accurately as possible. If a mistake was made, panellists would inform them by simply stating “incorrect, start again, 1687”. Panellists remained neutral throughout. For the control group, participants were given written maths problems to complete, without fear of evaluation. Once this task was over participants provided a fourth anxiety rating and second saliva sample before returning to room A. Following a rest period of 10 minutes a final anxiety rating and saliva sample was taken before participants were debriefed. The participant was reassured that no stress task would occur in the following days session, and that they would simply be completing more cognitive tasks and questionnaires. No mention was made of a follow up memory task. The following day participants completed the delayed memory task, which used same procedure as the immediate task. Participants then completed the PSRS before being fully debriefed as to the purposes of the experiment.

#### 2.4.2 Experiment 4: Stress at retrieval

The timeline for experiment 4 can be seen in Figure 1b below. The procedures were the same as in the previous experiments except the order in which they occurred was changed. The stress/control tasks were shifted in this experiment to the beginning of the second session, in day 2, after which the delayed retrieval was carried out.

### 2.5 Analysis of Stress Data

All participants provided three saliva samples at specific timepoints throughout the study (Figure 1). These samples were analysed in duplicate using commercially available cortisol ELISA kits (Salimetrics, UK). Single estimates of acute stress reactivity were measured as change in selfreported anxiety (sampled five times) and cortisol (from saliva sampled 3 times) were calculated using an area-under-the-curve analysis. Independent samples T-Tests were used to compare these measures of stress reactivity between stress and control groups.

To assess the relationship between these two measures of acute stress, a Pearson’s R correlation was used, based on all available data from all four experiments to provide a large enough sample to detect a medium effect size of .3 and with statistical power of .8 (minimum n required = 82) (Kirschbaum, Bartussek, & Strasburger, 1992). Including these correlations in each experiment separately would mean sample sizes are very small and would therefore be unstable estimates (Schönbrodt & Perugini, 2013). We also assessed the relationships between the two measures of acute stress (self-reported-anxiety and cortisol), and the relationship of each of these measures of acute stress separately with trait stress reactivity (measured using the PSRS). This correlation was run for stress and control groups separately, as the control group did not experience acute stress induction in the same way as the stress group did.

### 2.6 Analysis of Memory Data

Recognition accuracy was computed by converting the raw data into d’ scores, using standard procedures (Heeger, 1998). This analysis uses signal detection theory to account for measurement noise. Recollection scores were calculated using a corrected remember method by subtracting the false alarms (incorrectly endorsing new items as ‘old’) from hits (endorsing old items as old). Familiarity scores were calculated using the independent K method to establish familiarity scores that are unaffected by recollection scores (Yonelinas, 2002). We then computed persistence scores for each of these three measures, by dividing the scores for recognition accuracy, recollection, and familiarity the participant obtained on day 2 by the score they obtained on day 1 (Day 2/Day 1) (Craig, Della Sala, & Dewar, 2014). We opted to use persistence scores in favour of including delay as an additional independent variable to remove any group differences for baseline memory scores and acknowledge the non-linear nature of forgetting. Study 2 also used proportion scores for word pairs recalled during the cued recall part of the task, relative to the number of old responses correctly made. These scores were then also converted to a persistence score across days.

Potential outliers in persistence scores were identified using boxplots. Any data points scoring more than 3 interquartile ranges beyond the lower or upper quartile will be identified as extreme outliers and removed (Tukey, 1977).

The key analyses compared persistence scores between the stress and the control group. In Experiments 1, 2 and 4 the analysis used independent-samples T-test. In experiment 3, because there were three types of stimuli, we used mixed ANOVAs with group as a between-subject factor and list as a within-subject factor. Post hoc pairwise comparisons were then used to determine significant differences between specific list types.

Finally, memory persistence scores were correlated with stress scores (both self-reported anxiety and salivary concentrations) within the stress group only to assess if any relationship exists between the level of stress experiences and persistence rates. As the control group experienced little to no stress, we omitted them from this analysis.

## 3. Results common to all 4 experiments

We begin by reporting results for the stress manipulation, which was common to all 4 studies. Time course plots for self-reported anxiety and salivary cortisol can been seen in Figure 2. The independent T-test showed a significant difference between the stress and control groups for self-reported anxiety (*t*(142) = 6.10, *p* < .001). Similar significant differences are also seen between stress and control groups for salivary cortisol concentrations (*t*(85.489) = 3.33, *p* = .001). Descriptive statistics for measures of stress between groups shown in table 4.

We found a significant positive correlation between self-reported anxiety and salivary cortisol (r = .2, p = .025). Significant correlations between self-reported anxiety and trait stress were also seen in both the stress (*r* = .44, *p* < .001) and control (*r* = .26, *p* = .03) groups. Cortisol reactivity was not associated with trait stress in either stress (r = −.05, *p* = .71) or control (r = −.11, *p* = .38) groups.

**Figure 2.**
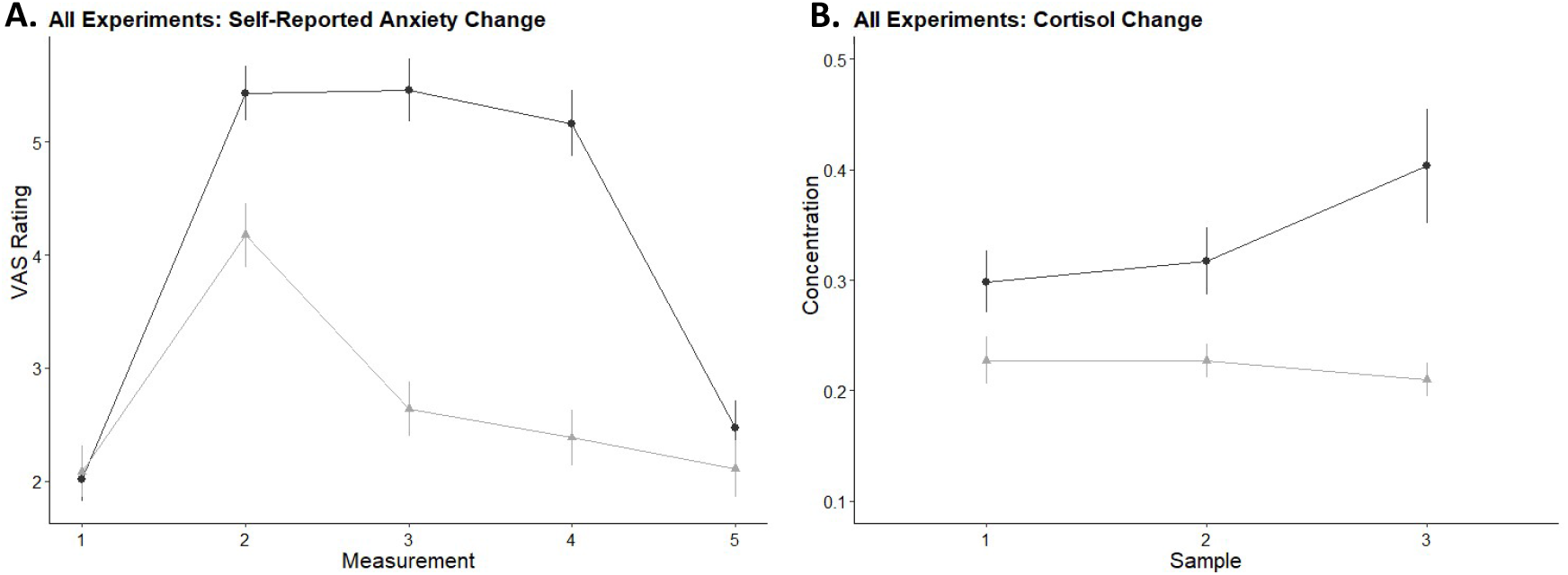
Error bars represent standard error. A) Mean self-report scores of anxiety for both groups at each of the five time points throughout the stress session: (1) immediately after baseline rest period (2) after preparation for the Trier (3) after the Trier speech task (4) after the Trier maths task (5) immediately before study debrief. B) Mean salivary cortisol concentrations measured at timepoints 1, 4 and 5.

**Table 4.** Mean and SD values for stress and control groups for measures of stress collapsed across all four experiments.

|  | Stress |  | Control |  |
| --- | --- | --- | --- | --- |
|  | Mean | SD | Mean | SD |
| Cortisol | .87 | .69 | .55 | .30 |
| Self-reported Anxiety | 20.02 | 6.97 | 12.95 | 6.94 |

### 4. Results from Experiment 1: Post encoding stress and neutral item recognition

#### 4.2.1 Measures of stress

The stress group reported higher levels of anxiety and showed higher cortisol reactivity than the control group (anxiety: *t*(34) = 3.76, *p* = .001. Cortisol: *t*(20.14) = 2.87, *p* = .01). Means and SD shown in table 5. Time course plots for self-reported anxiety and salivary cortisol can been seen below in figure 3.

**Table 5.** Experiment 1 group mean and SD’s for stress and memory measures.

|  | Stress |  | Control |  |
| --- | --- | --- | --- | --- |
|  | Mean | SD | Mean | SD |
| Cortisol | 1.4 | 0.9 | 0.7 | 0.4 |
| Self-reported Anxiety | 20.6 | 4.8 | 12.7 | 7.6 |
| Recognition | 0.13 | 0.71 | -0.11 | 0.64 |
| Recollection | 0.35 | 0.75 | 0.38 | 0.61 |
| Familiarity | 0.43 | 0.65 | 0.14 | 0.59 |

**Figure 3.**
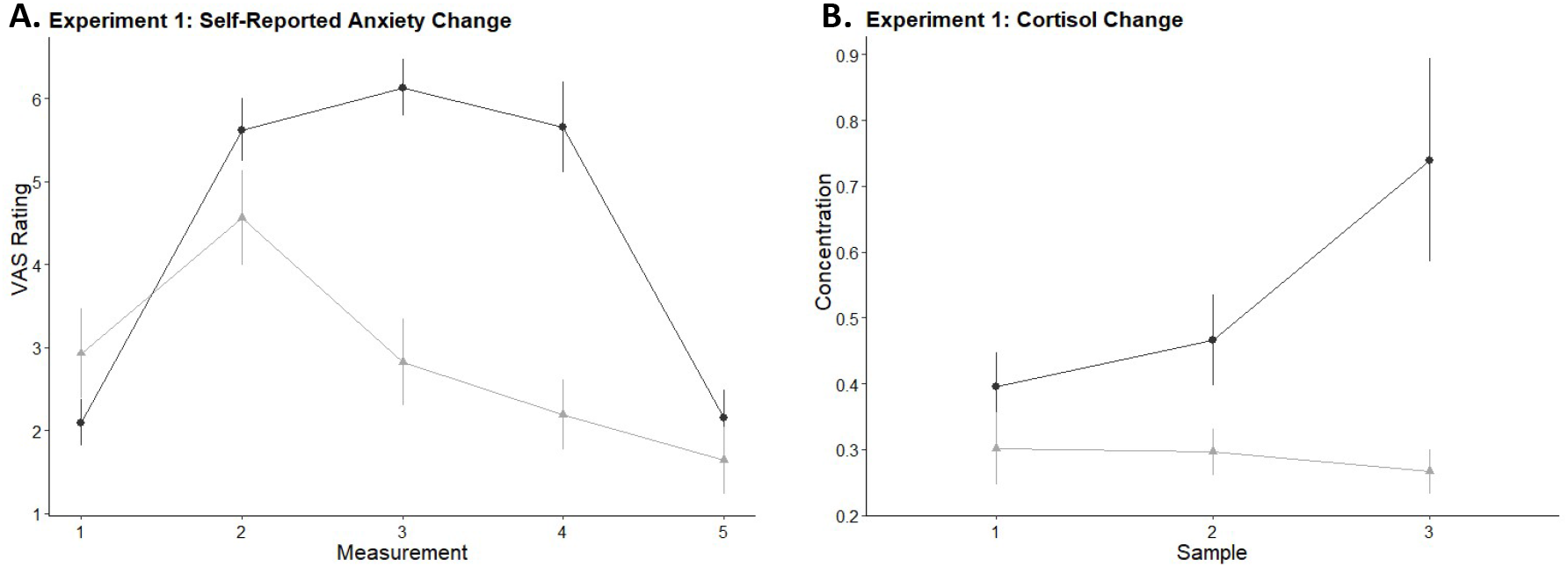
Error bars represent standard error. A) Mean self-report scores of anxiety for both groups at each of the five time points throughout the stress session: (1) immediately after baseline rest period (2) after preparation for the Trier (3) after the Trier speech task (4) after the Trier maths task (5) immediately before study debrief. B) Mean salivary cortisol concentrations measured at timepoints 1, 4 and 5.

No group differences were seen for scores of trait stress reactivity (p > .5) confirming that no prior differences exist between the groups for susceptibility to stress.

#### 4.2.2 Stress and memory

For overall recognition accuracy, no significant difference was shown between stress and control groups t(34)= 1.07, p = .294. Similarly, no group differences were observed between stress and controls for recollection t(34)=−.15, p = .879. Familiarity scores also did not differ between groups t(34) = 1.30, p = .2. Results are shown in Figure 4 and descriptive statistics in table 5.

**Figure 4.**
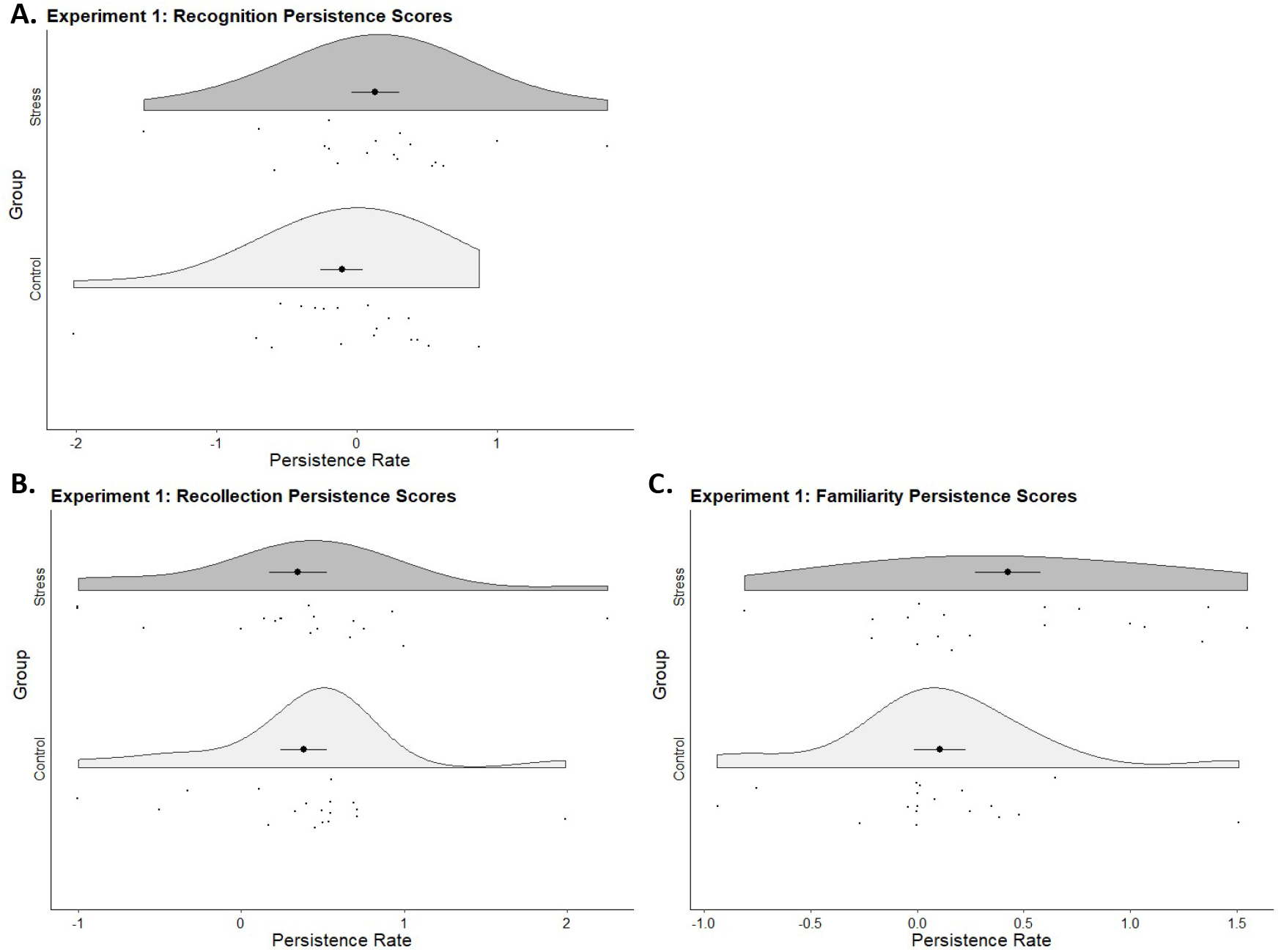
Raincloud plots to demonstrate the overall spread and individual data points representing persistence rates for recognition accuracy (A), recollection (B) and familiarity (C) for both stress and control groups. Bold central points with error bars represent mean persistence rates and standard error.

No significant correlations were seen between persistence scores for recognition accuracy or familiarity with either self-reported anxiety or cortisol. Recollection also did not significantly correlate with self-reported anxiety but show a strong significant positive correlation with cortisol response r = .66, p = .006.

### 5. Experiment 2. Post encoding stress and neutral association memory

#### 5.2.1 Measures of stress

Elevated levels of self-reported-anxiety were seen for participants in the stress group compared to the control group (*t*(1,34)= 2.5, *p* = .018; Figure 5). Salivary cortisol concentrations did not differ between stress and control groups during the stress task (*t*(1, 26)= 1.236, *p* = .227). Descriptive statistics shown in table 6.

**Figure 5.**
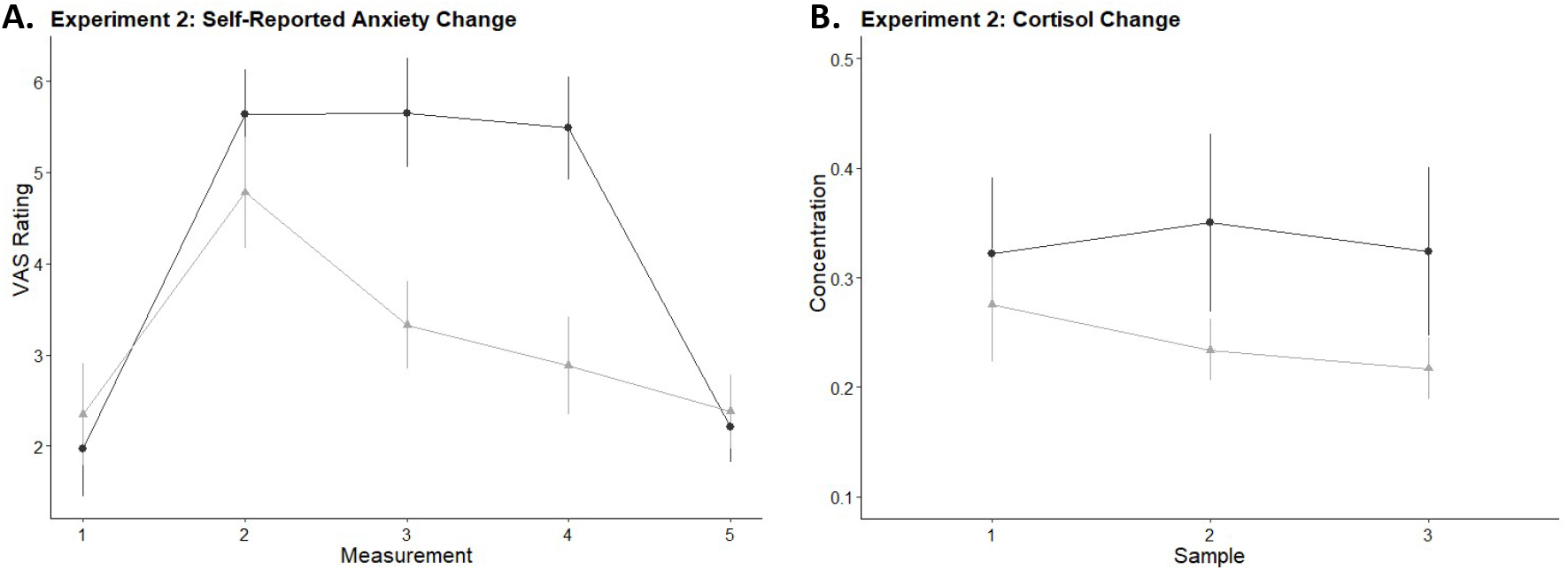
Error bars represent standard error. A) Mean self-report scores of anxiety for both groups at each of the five time points throughout the stress session: (1) immediately after baseline rest period (2) after preparation for the Trier (3) after the Trier speech task (4) after the Trier maths task (5) immediately before study debrief. B) Mean salivary cortisol concentrations measured at timepoints 1, 4 and 5.

**Table 6.**
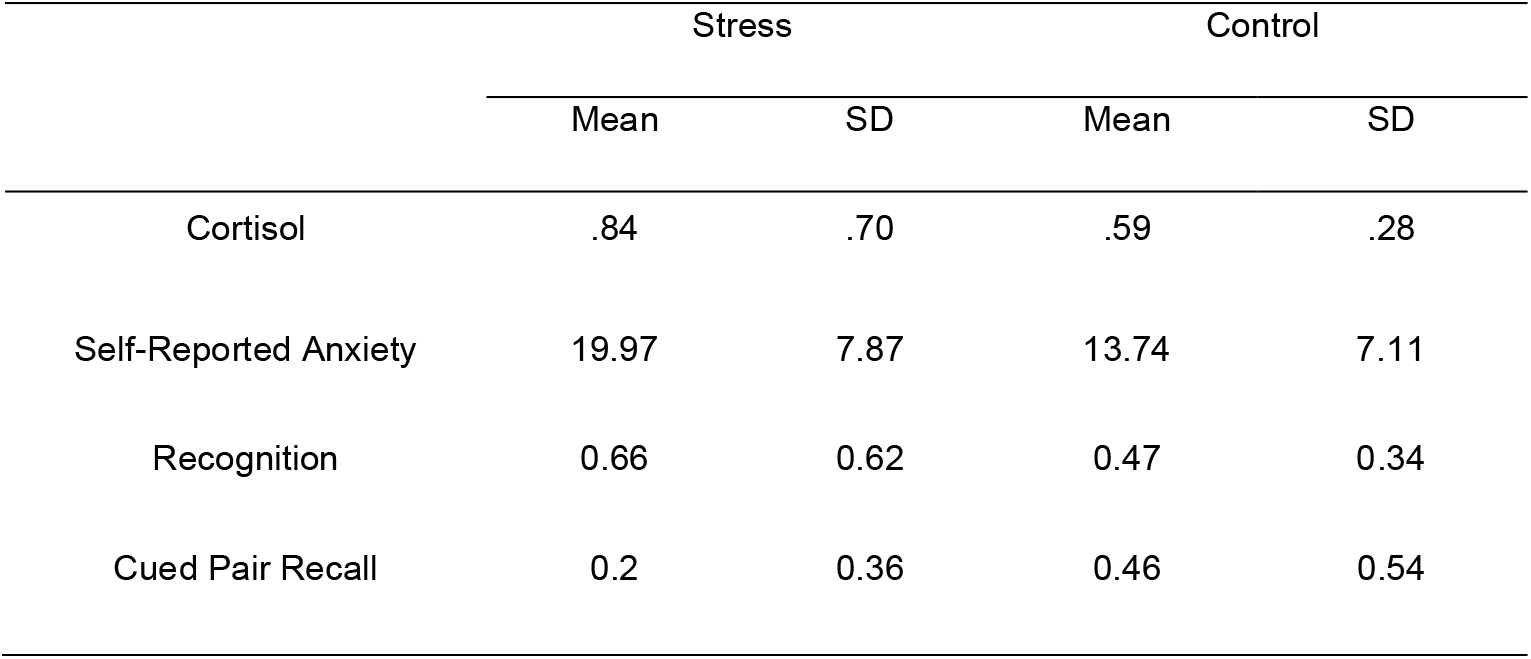
Experiment 2 group mean and SD’s for stress and memory measures.

|  | Stress |  | Control |  |
| --- | --- | --- | --- | --- |
|  | Mean | SD | Mean | SD |
| Cortisol | .84 | .70 | .59 | .28 |
| Self-Reported Anxiety | 19.97 | 7.87 | 13.74 | 7.11 |
| Recognition | 0.66 | 0.62 | 0.47 | 0.34 |
| Cued Pair Recall | 0.2 | 0.36 | 0.46 | 0.54 |

#### 5.2.2 Stress and memory

Independent samples t-tests were used to compare recognition scores between groups. Two participants’ recognition persistence scores were classified as outliers and removed. Additionally, one participant’s cued pair recall persistence score was identified as an outlier and removed. Removing these outliers did not affect the results of this experiment. Group means and SD shown in table 6. No significant group differences were seen between recognition accuracy scores *t*(27.045)= 1.151, *p* = .26 in the stress and control groups. No significant differences were seen between stress and control groups in cued pair recall persistence scores, *t*(29.887)= −1.68, *p* = .1. Recognition and cued recall persistence scores for both stress and control groups can be seen in figure 6. No significant correlations were seen between either memory or stress metric during this study.

**Figure 6.**
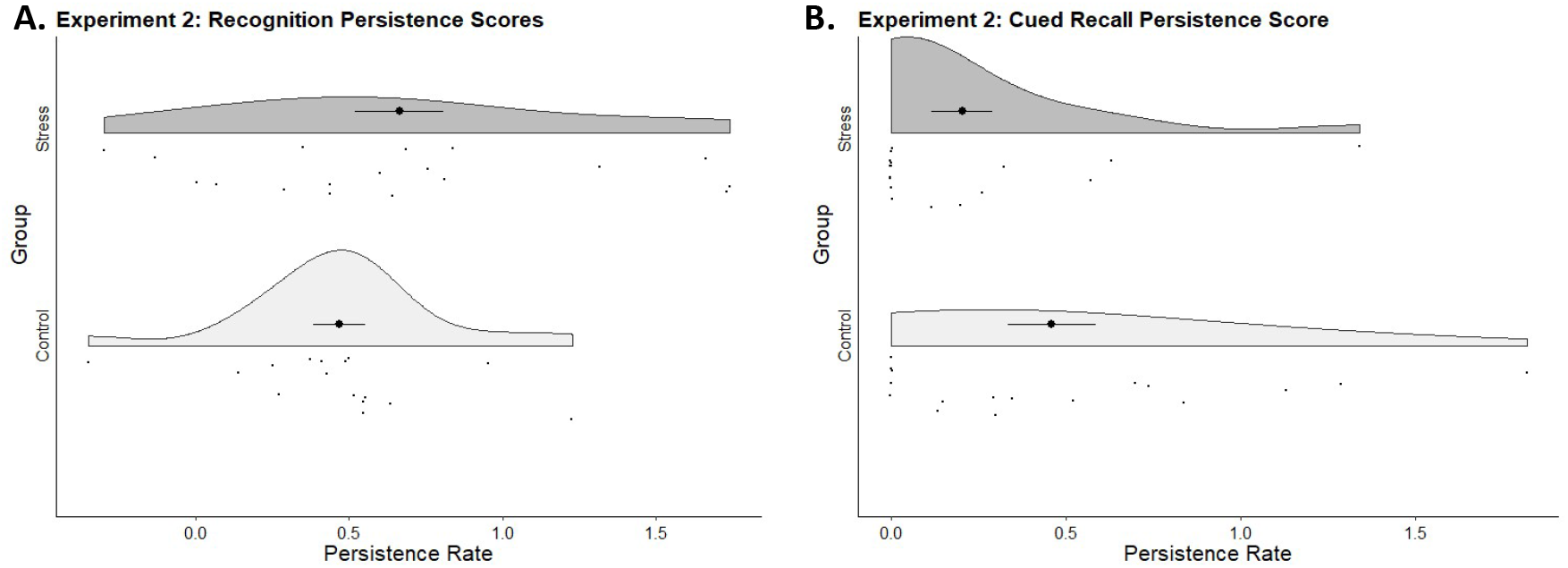
Raincloud plots to demonstrate the overall spread and individual data points representing persistence rates for recognition accuracy (A) and cued pair recall (B) in both stress and control groups. Bold central points with error bars represent mean persistence rates and standard error.

### 6. Experiment 3. Post encoding stress and emotional memory

#### 6.2.1 Measures of stress

The stress (self-reported anxiety: *M* = 19.80, *SD* = 6..98, cortisol: *M* = 0.72, *SD* = 0.48) group experienced greater acute stress reactivity than controls (self-reported anxiety: *M* = 11.13, *SD* = 6.35, cortisol: *M* = 0.41, *SD* = 0.16) both in terms of self-reported anxiety (*t*(34)= 3.9, *p* < .001) and cortisol, (*t*(28)=2.342, p = .027), see figure 7. Groups did not differ for trait stress reactivity (p > .45) confirming that no prior differences exist between the groups for self-reported susceptibility to stress.

**Figure 7.**
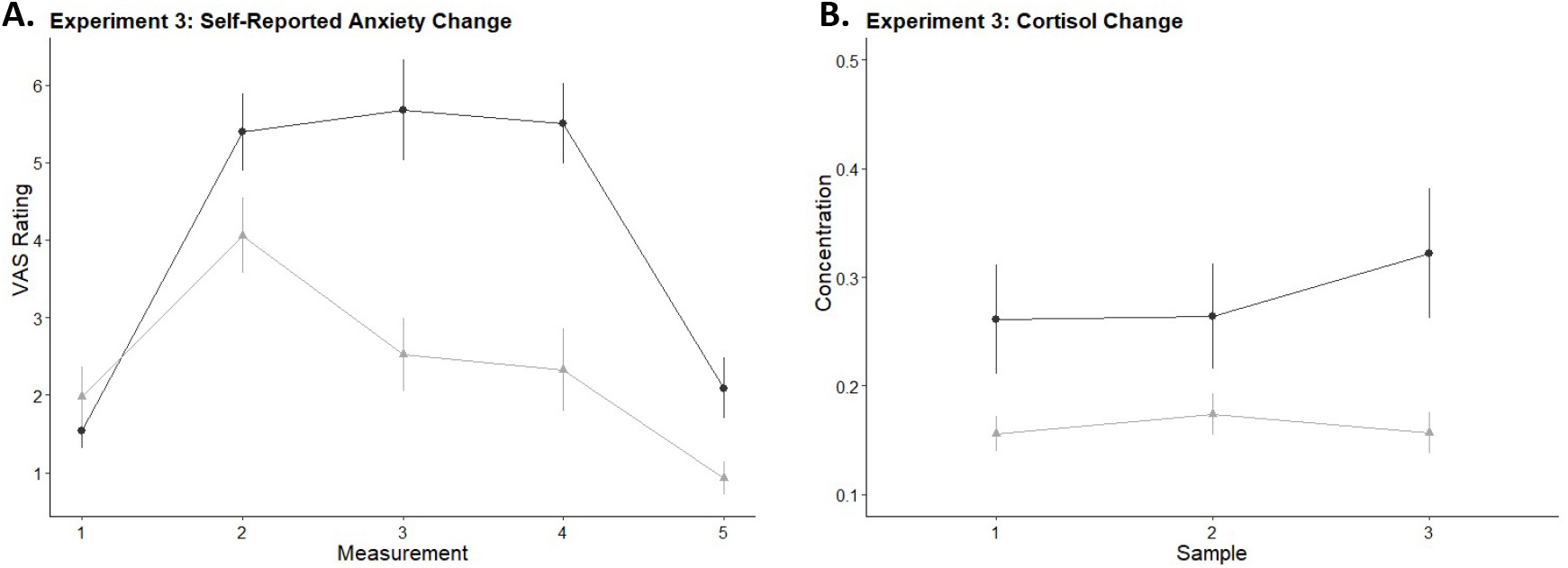
Error bars represent standard error. A) Mean self-report scores of anxiety for both groups at each of the five time points throughout the stress session: (1) immediately after baseline rest period (2) after preparation for the Trier (3) after the Trier speech task (4) after the Trier maths task (5) immediately before study debrief. B) Mean salivary cortisol concentrations measured at timepoints 1, 4 and 5.

#### 6.2.2 Stress and emotional memory

Persistence scores for recognition accuracy, recollection and familiarity were compared using mixed 2 (group) x 3 (list type) ANOVAs. Descriptive statistics for groups overall and separated by list type are shown in table 7. For recognition accuracy, no significant effects of list type or group were observed, neither did the two factors interact. No significant main effects or interactions were found between stress and control groups for familiarity scores. For recollection persistence rates however, a significant list type x group interaction was seen (F(2, 62)=3.7, p = .03), in the absence of main effects of either factor. Figure 8 shows stress and control group persistence scores for each memory metric.

**Table 7.** Experiment 3 memory metric group means and SD below in brackets for each memory metric, split by list type.

|  | Negative-Emotional |  | Related-Neutral |  | Random-Neutral |  |
| --- | --- | --- | --- | --- | --- | --- |
|  | Stress | Control | Stress | Control | Stress | Control |
| Recognition | 0.48 | 0.61 | 0.51 | 0.56 | 0.62 | 0.59 |
| accuracy | (0.27) | (0.27) | (0.26) | (0.53) | (0.33) | (0.42) |
| Recollection | 0.57 | 0.72 | 0.78 | 0.40 | 0.57 | 0.69 |
|  | (0.52) | (0.32) | (0.46) | (0.34) | (0.57) | (0.49) |
| Familiarity | 2.04 | -1.73 | -3.83 | -0.11 | -0.36 | -7.28 |
|  | (18.75) | (6.33) | (23.03) | (1.10) | (1.85) | (24.43) |

**Figure 8.**
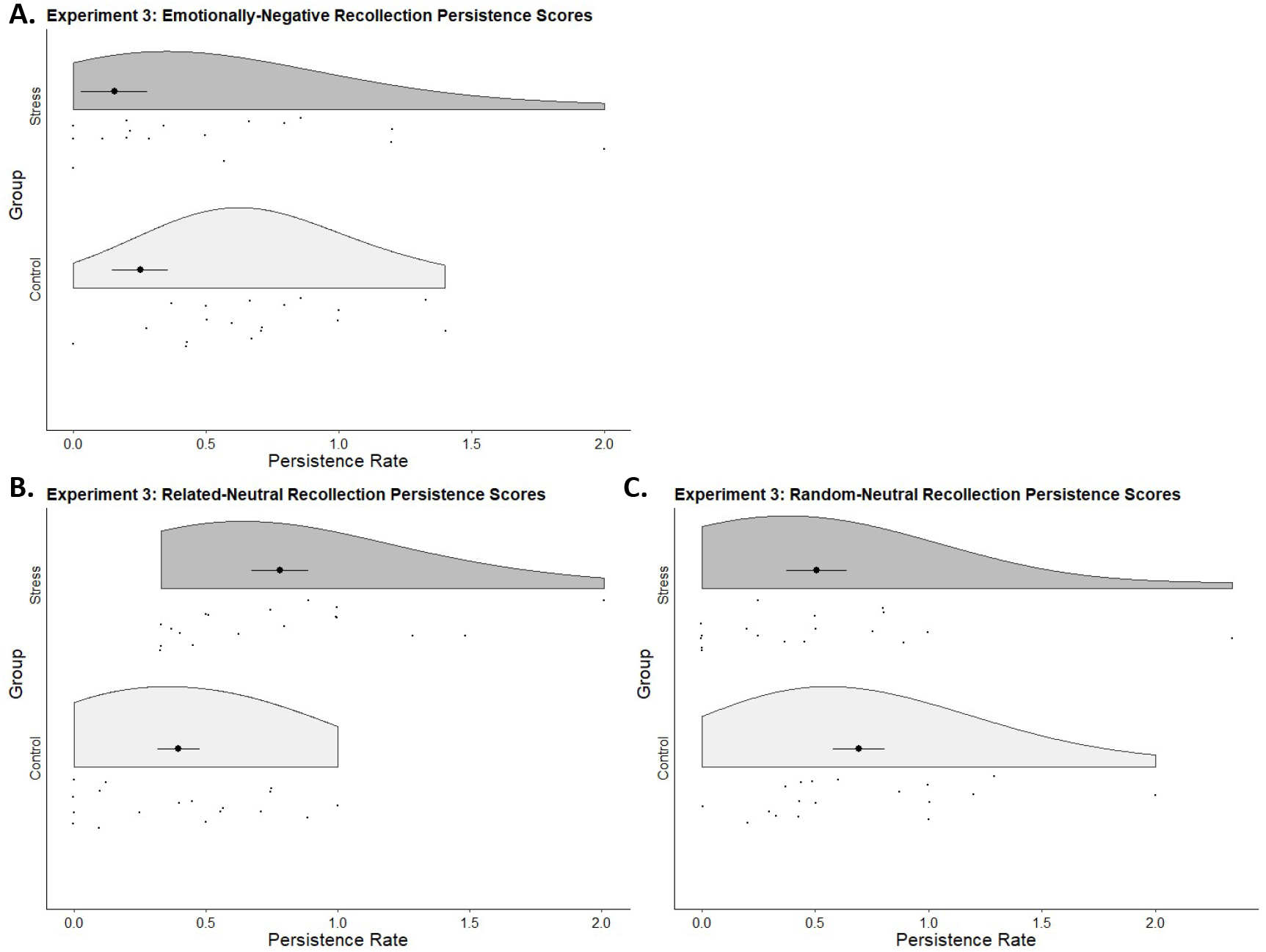
Raincloud plots to demonstrate the overall spread and individual data points representing persistence rates of recollection for emotionally-negative (A), related-neutral (B) and random-neutral (C) stimuli in stress and control groups. Bold central points with error bars represent mean persistence rates and standard error.

**Figure 9.**
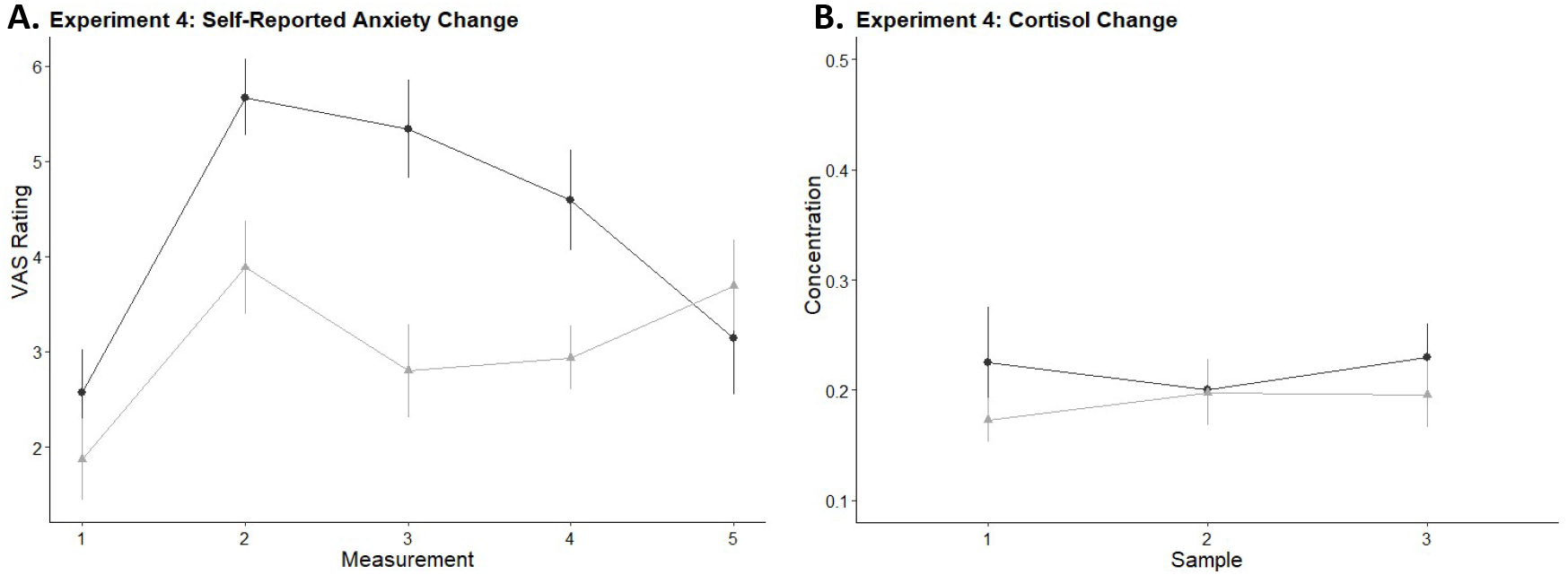
Error bars represent standard error. A) Mean self-report scores of anxiety for both groups at each of the five time points throughout the stress session: (1) immediately after baseline rest period (2) after preparation for the Trier (3) after the Trier speech task (4) after the Trier maths task (5) immediately before study debrief. B) Mean salivary cortisol concentrations measured at timepoints 1, 4 and 5.

**Figure 10.**
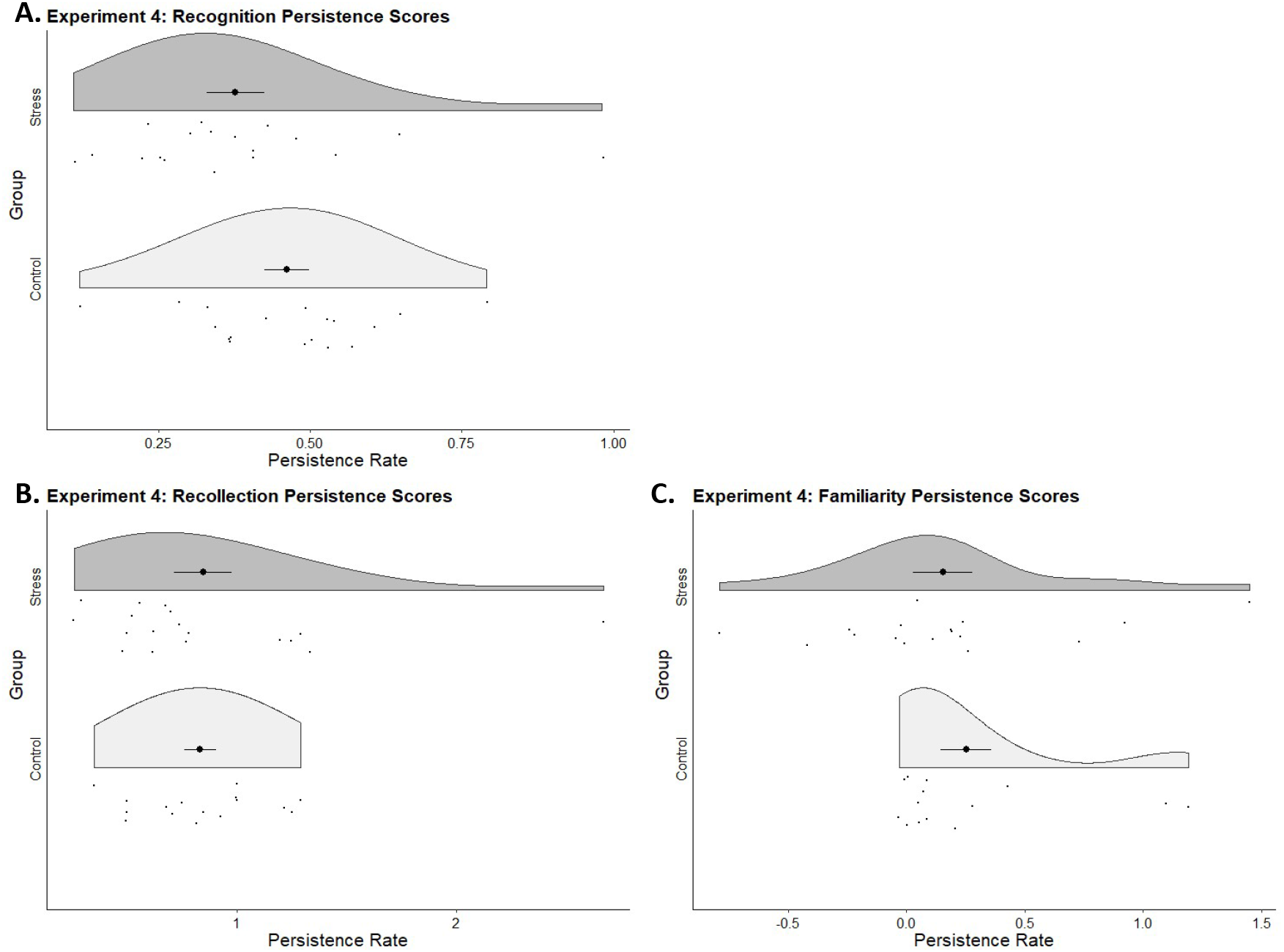
Raincloud plots to demonstrate the overall spread and individual data points representing persistence rates for recognition accuracy (A), recollection (B) and familiarity (C) for both stress and control groups. Bold central points with error bars represent mean persistence rates and standard error.

Post-hoc analyses were used to explore the significant interaction between list type and group for recollection persistence rates. Analyses were conducted separately for list type to compare persistence scores for each group. For emotionally-negative stimuli and random-neutral stimuli, stress and control groups did not differ in terms of recollection persistence scores (*t*(32)=−1.04, p = .31 and *t*(32)=−0.68, p = .5 respectively). Persistence scores for related-neutral items however, were significantly reduced for the control group compared to stress, *t*(34)= 2.86, p = .007.

No significant relationship was shown between any stress, list type or memory measure. There was, however, a significant positive correlation between self-reported anxiety and recollection responses, *r* = 4.96, p = .036.

### 7. Experiment 4. Stress at retrieval

#### 7.2.1 Measures of stress

Independent samples t-tests confirmed that overall, the stress group reported significantly higher levels of anxiety during the task than the control group (*t*(34) = 2.16, p = .038), but did not exhibit a greater cortisol response (*t*(32)=.71, *p* = .48 (Figure 11). No group differences were seen for trait stress reactivity (p > .25) confirming that no prior differences exist between the groups for selfreported susceptibility to stress. All descriptive statistics are shown in table 8.

**Table 8.** Experiment 4 group mean and SD’s for stress and memory measures.

|  | Stress |  | Control |  |
| --- | --- | --- | --- | --- |
|  | Mean | SD | Mean | SD |
| Cortisol | 0.55 | 0.28 | 0.48 | 0.27 |
| Self-reported Anxiety | 19.72 | 8.27 | 14.25 | 6.88 |
| Recognition | 0.38 | 0.2 | 0.46 | 0.15 |
| Recollection | 0.85 | 0.56 | 0.84 | 0.29 |
| Familiarity | 0.16 | 0.51 | 0.25 | 0.4 |

#### 7.2.2 Stress and memory

Independent samples t-tests were used to compare groups’ persistence scores for recognition accuracy, recollection and familiarity. Two recollection persistence scores and five familiarity persistence scores were identified as outliers and removed. The results of this experiment were not affected by removing outliers. No significant differences were seen between stress and control groups for recognition accuracy (*t*(34)= −1.42, *p* = .16). Similarly, no differences were seen between groups for recollection scores (*t*(32)=0.08, *p* = .94) or between groups for familiarity scores (*t*(29)=−0.58, *p* = .57). No significant correlations are seen between either memory or stress metric during this study.

## 8. Discussion

The current set of experiments cautiously and carefully assessed the effects of stress on episodic memory. Using a stepped approach, we provide reproducible evidence that our psychosocial stress test significantly increased perceived anxiety and salivary cortisol levels (when considered overall) but had a limited effect on recognition memory accuracy, recollection, familiarity, and on associative memory, measured through cued recall. In particular, we found no significant effects of psychosocial stress, induced either after learning or before retrieval, on item recognition or on cued-recall. Participants who experienced greater stress did, however, show greater change in recollection in some experiments. Additionally, the stress group in experiment 3 showed more persistence in their recollection of related-neutral items compared to controls. Together these findings suggest that the impact of psychosocial stress on memory has been overstated in the literature but that inter-individual differences may help to explain some of the discrepancies in findings, particularly when recollection is assessed.

In line with previous social stress studies (Corbett et al., 2017), we found no significant effects of experimentally induced psychosocial stress on recognition accuracy or familiarity. This was seen in experiments one, two and three. In each of these experiments, we see no difference in persistence rates between stress and control groups. This was extended in experiment 4, where we also found no significant effect of stress immediately before retrieval on recognition accuracy or familiarity. These findings differ from studies using physical stressors, which often report that postencoding stress enhances recognition memory, in particular familiarity (McCullough & Yonelinas, 2013; Yonelinas, Parks, Koen, Jorgenson, & Mendoza, 2011). A potential reason for this difference in findings may be that physiological stress is associated with greater activation of the temporal pole (Kogler et al., 2015), and these anterior temporal regions are more closely associated with familiarity memory (Yonelinas, Otten, Shaw, & Rugg, 2005). Together with our own results, these findings challenge the generally upheld view that all forms of post-encoding stress enhance all forms of memory, while stress at retrieval unanimously impairs it (Shields et al., 2017; Wolf, 2009).

We did, however, observe some significant positive correlations for stress and recollection in experiment 1 for cortisol and experiment 3 for self-reported anxiety. These findings indicate a relationship between level of stress experienced and recollection ability within the stress group. Given that these associations were observed on different measures of stress (self-perceived vs cortisol), and that such correlations within each experiment were likely underpowered, we should be careful in how we interpret these findings. But they do suggest a relationship between high levels of post-encoding stress and enhanced recollection. If replicated, these findings would confirm that each individual’s level of stress reactivity and subsequent experience of the stressor, may be what dictates the effects of stress on cognition (Nater et al., 2007; Oei, Everaerd, Elzinga, Van Well, & Bermond, 2006; Takahashi et al., 2004). To better explore this effect, future research could design and power studies appropriately to be able to better compare responders and non-responders to stress, be it pre-defining participants as high and low stress reactive or defining groups post-hoc based on their stress response.

In experiment three, recollection of related-neutral words showed greater rates of persistence in the stress group than in controls. This finding suggests that social stress specifically enhances recollection persistence for words that are highly semantically-related. This could be expanded to explain why stress has previously been thought to increase memory for emotional over neutral stimuli (Buchanan & Lovallo, 2001; Jelici, Geraerts, Merckelbach, & Guerrieri, 2004), as emotional stimuli is thought to be more intrinsically semantically related than neutral stimuli (Buchanan, Etzel, Adolphs, & Tranel, 2006; Talmi & Moscovitch, 2004). In our case, related-neutral stimuli were more semantically-related than both emotional and random-neutral words, as shown by significant differences in LSA scores between list types. It could therefore be suggested that psychosocial stress only modulates recollection of words that are highly semantically-related, regardless of their emotional valence. Future studies should carefully consider the semantic relatedness of their emotional (as well as their neutral) stimuli.

In addition to the stepped replication design used for these experiments, a further benefit of this study is that unlike many others, we also examine individuals’ trait stress reactivity and their subjective experience of stress during the TSST in addition to their cortisol reactivity. Self-reported measured of anxiety during the stress task provided an insight into emotional and cognitive responses to stress (Campbell & Ehlert, 2012), offering a quick and more holistic view of the individuals experience of the stress task than cortisol measures alone (Lazarus & Folkman, 1984; Schlotz et al., 2011). Although potentially susceptible to bias, self-reports of anxiety, measured using visual analogue scales, have been shown to be reliably sensitive to changes in mental state, such as those experienced during a social stress task (Cella & Perry, 1986; Taylor et al., 2018).

There were however some limitations that must be considered. For instance, although selfreported anxiety was seen to increase significantly in the stress over control groups, the rise in cortisol levels did not always reach significance in individual experiments. Increases in salivary cortisol that do not reach significance have been seen in other studies within the stress and cognition literature, both for psychosocial and physical stressors (Duncko, Johnson, Merikangas, & Grillon, 2009; Jelici et al., 2004). Critically, when pooling the findings across all four experiments we did see significant differences between stress and control groups for both cortisol levels and self-reports of anxiety, suggesting it may have been an issue of power when considering individual experiments alone. We also show an overall significant positive relationship between self-reported anxiety and in cortisol, suggesting that the stress manipulation had the desired effects.

A further potential limitation is that our participants were exposed to only moderate levels of stress. More extreme stressful situations could have had more dramatic effects on memory. Alternatively, we could stipulate a minimum level of stress experienced for each individual participant, this would help to minimise the influence of inter-individual differences in stress-reactivity. This suggestion is supported by the significant relationship we see between recollection and stress experienced in experiments 1 & 3. Discovering where boundaries for this effect lie would help us understand exactly how different forms of stress affect episodic memory.

In summary, we provide strong evidence to demonstrate that mild to moderate levels of psychosocial stress has minimal effects on item recognition accuracy at the group level, regardless of the timing of stress. Instead, we demonstrate a potential relationship between greater levels of post encoding stress and enhanced subsequent recollection within the stress group. We also find no evidence to suggest any such relationship exists between levels of social stress and familiarity. We can speculate that generally greater responsivity to stress may be required to influence recollection. We therefore believe that individual response to such tasks is an important and relevant factor and future research should attempt to select experimental samples based on levels of stress-reactivity. Finally, we also observed that post-encoding stress enhances recollection persistence rates of semantically-related words relative to controls. Based on findings from both experiment 1 and 3, we propose that psychosocial stress acts upon hippocampally-dependent memory processes, such as recollection, particularly for more highly semantically-related material. As our findings consistently differ from reports of the impact of physical stress on memory, this may be critical to consider when explaining the variability in findings in the literature.

## Acknowledgements

This work was funded by the BBSRC as part of a DTP studentship. We would also like to thank and acknowledge the hard work of all the undergraduate students in the Muhlert lab (2017-2019) who have been involved in the data collection for all four of these experiments.

## Author contributions

All authors were involved in the formulation of the research ideas and development of methodology. Elizabeth McManus collected and analysed the data and created the original draft. Deborah Talmi, Hamied Haroon and Nils Muhlert provided critical reviews of the drafts and acquired financial support for the project.

## Abbreviations

TSST: Trier Social Stress Test

## References

Beckner, V. E., Tucker, D. M., Delville, Y., & Mohr, D. C. (2006). Stress facilitates consolidation of verbal memory for a film but does not affect retrieval. Behavioral Neuroscience, 120(3), 518–527. https://doi.org/10.1037/0735-7044.120.3.518

Buchanan, T. W., Etzel, J. A., Adolphs, R., & Tranel, D. (2006). The influence of autonomic arousal and semantic relatedness on memory for emotional words. International Journal of Psychophysiology, 61(1), 26–33.

Buchanan, T. W., & Lovallo, W. R. (2001). Enhanced memory for emotional material following stress-level cortisol treatment in humans. Psychoneuroendocrinology, 26(3), 307–317. https://doi.org/10.1016/S0306-4530(00)00058-5

Campbell, J., & Ehlert, U. (2012). Acute psychosocial stress: does the emotional stress response correspond with physiological responses? Psychoneuroendocrinology, 37(8), 1111–1134.

Cella, D. F., & Perry, S. W. (1986). Reliability and concurrent validity of three visual-analogue mood scales. Psychological Reports, 59(2), 827–833.

Chida, Y., & Hamer, M. (2008). Chronic psychosocial factors and acute physiological responses to laboratory-induced stress in healthy populations: a quantitative review of 30 years of investigations. Psychological Bulletin, 134(6), 829–885. https://doi.org/10.1037/a0013342

Cohen, J. (1988). The effect size. Statistical Power Analysis for the Behavioral Sciences, 77–83.

Corbett, B., Weinberg, L., & Duarte, A. (2017). The effect of mild acute stress during memory consolidation on emotional recognition memory. Neurobiology of Learning and Memory, 145, 34–44. https://doi.org/10.1016/j.nlm.2017.08.005

Cornelisse, S., van Stegeren, A. H., & Joëls, M. (2011). Implications of psychosocial stress on memory formation in a typical male versus female student sample. Psychoneuroendocrinology, 36(4), 569–578. https://doi.org/10.1016/j.psyneuen.2010.09.002

Craig, M., Della Sala, S., & Dewar, M. (2014). Autobiographical thinking interferes with episodic memory consolidation. PloS One, 9(4), e93915.

Dedovic, K., Duchesne, A., Andrews, J., Engert, V., & Pruessner, J. C. (2009). The brain and the stress axis: the neural correlates of cortisol regulation in response to stress. Neuroimage, 47(3), 864–871.

Duncko, R., Johnson, L., Merikangas, K., & Grillon, C. (2009). Working memory performance after acute exposure to the cold pressor stress in healthy volunteers. Neurobiology of Learning and Memory, 91(4), 377–381.

Elzinga, B. M., Bakker, A., & Bremner, J. D. (2005). Stress-induced cortisol elevations are associated with impaired delayed, but not immediate recall. Psychiatry Research, 134(3), 211–223. https://doi.org/10.1016/j.psychres.2004.11.007

Fries, E., Dettenborn, L., & Kirschbaum, C. (2009). The cortisol awakening response (CAR): facts and future directions. International Journal of Psychophysiology, 72(1), 67–73.

Grühn, D. (2016). An English Word Database of EMOtional TErms (EMOTE). Psychological Reports, 119(1), 290–308.

Heeger, D. (1998). Signal detection theory. California.

Jelici, M., Geraerts, E., Merckelbach, H., & Guerrieri, R. (2004). Acute stress enhances memory for emotional words, but impairs memory for neutral words. International Journal of Neuroscience, 114(10), 1343–1351.

Kamp, S., Endemann, R., Domes, G., & Mecklinger, A. (2019). Neurobiology of Learning and Memory E ff ects of acute psychosocial stress on the neural correlates of episodic encoding : Item versus associative memory. Neurobiology of Learning and Memory, 157(November 2018), 128–138. https://doi.org/10.1016/j.nlm.2018.12.006

Kirschbaum, C., Bartussek, D., & Strasburger, C. J. (1992). Cortisol responses to psychological stress and correlations with personality traits. Personality and Individual Differences, 13(12), 1353–1357.

Kirschbaum, C., Pirke, K.-M., & Hellhammer, D. H. (1993). The ‘Trier Social Stress Test’–a tool for investigating psychobiological stress responses in a laboratory setting. Neuropsychobiology, 28(1–2), 76–81.

Kogler, L., Müller, V. I., Chang, A., Eickhoff, S. B., Fox, P. T., Gur, R. C., & Derntl, B. (2015). Psychosocial versus physiological stress - Meta-analyses on deactivations and activations of the neural correlates of stress reactions. NeuroImage, 119, 235–251. https://doi.org/10.1016/j.neuroimage.2015.06.059

Kuhlmann, S., Piel, M., & Wolf, O. T. (2005). Impaired memory retrieval after psychosocial stress in healthy young men. Journal of Neuroscience, 25(11), 2977–2982.

LaBar, K. S., & Cabeza, R. (2006). Cognitive neuroscience of emotional memory. Nature Reviews Neuroscience, 7(1), 54.

Lazarus, R. S., & Folkman, S. (1984). Stress, appraisal, and coping. Springer publishing company.

Li, S., Weerda, R., Guenzel, F., Wolf, O. T., & Thiel, C. M. (2013). ADRA2B genotype modulates effects of acute psychosocial stress on emotional memory retrieval in healthy young men. Neurobiology of Learning and Memory, 103, 11–18.

Lupien, S. J., & Lepage, M. (2001). Stress, memory, and the hippocampus: Can’t live with it, can’t live without it. Behavioural Brain Research, 127(1–2), 137–158. https://doi.org/10.1016/S0166-4328(01)00361-8

McCullough, A. M., & Yonelinas, A. P. (2013). Cold-pressor stress after learning enhances familiarity-based recognition memory in men. Neurobiology of Learning and Memory, 106, 11–17.

McEwen, B. S. (2007). Physiology and neurobiology of stress and adaptation: central role of the brain. Physiological Reviews, 87(3), 873–904.

McGaugh, J. L. (2004). The amygdala modulates the consolidation of memories of emotionally arousing experiences. Annu. Rev. Neurosci., 27, 1–28.

Migo, E. M., Mayes, A. R., & Montaldi, D. (2012). Measuring recollection and familiarity: Improving the remember/know procedure. Consciousness and Cognition, 21(3), 1435–1455.

Montaldi, D., & Mayes, A. R. (2010). The role of recollection and familiarity in the functional differentiation of the medial temporal lobes. Hippocampus, 20(11), 1291–1314.

Nater, U. M., Moor, C., Okere, U., Stallkamp, R., Martin, M., Ehlert, U., & Kliegel, M. (2007). Performance on a declarative memory task is better in high than low cortisol responders to psychosocial stress. Psychoneuroendocrinology, 32(6), 758–763. https://doi.org/10.1016/j.psyneuen.2007.05.006

Oei, N. Y. L., Everaerd, W. T. A. M., Elzinga, B. M., Van Well, S., & Bermond, B. (2006). Psychosocial stress impairs working memory at high loads: An association with cortisol levels and memory retrieval. Stress, 9(3), 133–141. https://doi.org/10.1080/10253890600965773

Old, S. R., & Naveh-Benjamin, M. (2008). Differential effects of age on item and associative measures of memory: a meta-analysis. Psychology and Aging, 23(1), 104.

Ponzio, A., & Mather, M. (2014). Hearing something emotional influences memory for what was just seen: How arousal amplifies effects of competition in memory consolidation. Emotion, 14(6), 1137.

Preuß, D., & Wolf, O. T. (2009). Neurobiology of Learning and Memory Post-learning psychosocial stress enhances consolidation of neutral stimuli. Neurobiology of Learning and Memory, 92(3), 318–326. https://doi.org/10.1016/j.nlm.2009.03.009

Schlotz, W., Yim, I. S., Zoccola, P. M., Jansen, L., & Schulz, P. (2011). The Perceived Stress Reactivity Scale: Measurement Invariance, Stability, and Validity in Three Countries. Psychological Assessment, 23(1), 80–94. https://doi.org/10.1037/a0021148

Schönbrodt, F., & Perugini, M. (2013). At what sample size do correlations stabilize? Journal of Research in Personality, 47. https://doi.org/10.1016/j.jrp.2013.05.009

Schoofs, D., & Wolf, O. T. (2009). Stress and Memory Retrieval in Women: No Strong Impairing Effect During the Luteal Phase. Behavioral Neuroscience, 123(3), 547–554. https://doi.org/10.1037/a0015625

Sharot, T., & Yonelinas, A. P. (2008). Differential time-dependent effects of emotion on recollective experience and memory for contextual information. Cognition, 106(1), 538–547.

Sheldon, S., Chu, S., Nitschke, J. P., Pruessner, J. C., & Bartz, J. A. (2018). The dynamic interplay between acute psychosocial stress, emotion and autobiographical memory. Scientific Reports, (May), 1–12. https://doi.org/10.1038/s41598-018-26890-8

Shields, G. S., Sazma, M. A., McCullough, A. M., & Yonelinas, A. P. (2017). The effects of acute stress on episodic memory: A meta-analysis and integrative review. Psychological Bulletin, 143(6), 636.

Takahashi, T., Ikeda, K., Ishikawa, M., Tsukasaki, T., Nakama, D., Tanida, S., & Kameda, T. (2004). Social stress-induced cortisol elevation acutely impairs social memory in humans. Neuroscience Letters, 363(2), 125–130. https://doi.org/10.1016/j.neulet.2004.03.062

Talmi, D., & Moscovitch, M. (2004). Can semantic relatedness explain the enhancement of memory for emotional words? Memory and Cognition, 32(5), 742–751. https://doi.org/10.3758/BF03195864

Taylor, J. H., Landeros-Weisenberger, A., Coughlin, C., Mulqueen, J., Johnson, J. A., Gabriel, D., … Bloch, M. H. (2018). Ketamine for social anxiety disorder: A randomized, placebo-controlled crossover trial. Neuropsychopharmacology, 43(2), 325–333.

Tollenaar, M. S., Elzinga, B. M., Spinhoven, P., & Everaerd, W. (2009). Autobiographical memory after acute stress in healthy young men. Memory, 17(3), 301–310. https://doi.org/10.1080/09658210802665845

Tukey, J. W. (1977). Exploratory data analysis (Vol. 2). Reading, MA.

Wolf, O. T. (2009). Stress and memory in humans: twelve years of progress? Brain Research, 1293, 142–154.

Wolf, O. T. (2012). Immediate recall influences the effects of pre-encoding stress on emotional episodic long-term memory consolidation in healthy young men. Stress, 15(3), 272–280. https://doi.org/10.3109/10253890.2011.622012

Wolf, O. T. (2017). Stress and memory retrieval: mechanisms and consequences. Current Opinion in Behavioral Sciences, 14, 40–46.

Yonelinas, A. P. (2002). The nature of recollection and familiarity: A review of 30 years of research. Journal of Memory and Language, 46(3), 441–517.

Yonelinas, A. P., Otten, L. J., Shaw, K. N., & Rugg, M. D. (2005). Separating the brain regions involved in recollection and familiarity in recognition memory. Journal of Neuroscience, 25(11), 3002–3008.

Yonelinas, A. P., Parks, C. M., Koen, J. D., Jorgenson, J., & Mendoza, S. P. (2011). The effects of postencoding stress on recognition memory: examining the impact of skydiving in young men and women. Stress, 14(2), 136–144.

